# MELD: Mixed Effects for Large Datasets

**DOI:** 10.1101/156315

**Authors:** Dylan M. Nielson, Per B. Sederberg

## Abstract

Mixed effects models provide significant advantages in sensitivity and flexibility over typical statistical approaches to neural data analysis, but mass univariate application of mixed effects models to large neural datasets is computationally intensive. Threshold free cluster enhancement also provides a significant increase in sensitivity, but requires computationally-intensive permutation-based significance testing. Not surprisingly, the combination of mixed effects models with threshold free cluster enhancement and nonparametric permutation-based significance testing is currently completely impractical. With mixed effects for large datasets (MELD) we circumvent this impasse by means of a singular value decomposition to reduce the dimensionality of neural data while maximizing signal. Singular value decompositions become unstable when there are large numbers of noise features, so we precede it with a bootstrap-based feature selection step employing threshold free cluster enhancement to identify stable features across subjects. By projecting the dependent data into the reduced space of the singular value decomposition we gain the power of a multivariate approach and we can greatly reduce the number of mixed effects models that need to be run, making it feasible to use permutation testing to determine feature level significance. Due to these innovations, MELD is much faster than an element-wise mixed effects analysis, and on simulated data MELD was more sensitive than standard techniques, such as element-wise t-tests combined with threshold-free cluster enhancement. When evaluated on an EEG dataset, MELD identified more significant features than the t-tests with threshold free cluster enhancement in a comparable amount of time.

## Introduction

The design of most neuroimaging research, and especially electroencephalography (EEG) or magnetoencephalography (MEG), seeks to identify the subset of sensors and times after stimulus onset where different classes of trials elicit distinct neural signals. In EEG and MEG the most common way to analyze these experiments is to average neural responses for different classes of trials within participants and then to perform an across subject ANOVA or t-test comparing locations and time points selected a priori (1,2). Nonparametric permutation testing can provide a means to account for multiple comparisons, allowing all sensors and time points to be tested (3). Furthermore, employing threshold free cluster enhancement (TFCE) of the test statistic (e.g., *t* values) to account for autocorrelation between spatiotemporal features in neural data significantly improves the sensitivity of this approach (4–6). The power and simplicity of nonparametric t-tests or ANOVAs with TFCE have led to its widespread use in neuroimaging and electrophysiology, but the reliance of this approach on within-subject averages and simple regression limits experimental designs and is unable to account for known sources of trial level variance, reducing its sensitivity.

The standard approach assumes that within each subject there is a consistent or “fixed” effect driving the difference between classes of trials and that any trial level variance can be neglected because it affects both classes of trials equally. The within subject effect is assessed by the first-level test, which may be as simple as a mean difference, or it may be a more complex model such as a GLM. The second-level test then assesses the significance of the subject level effects only taking into account variance across subjects. If there are known confounds at the trial level, they must be accounted for via counterbalancing, significantly complicating experimental design. Mixed effects models, such as those implemented in linear mixed effects regression (LMER), avoid these issues by simultaneously modeling fixed effects and “random” effects (sources of variance) at multiple levels (7–9). Mixed effects models are well established in linguistics research, and their utility for event related potential (ERP) analysis has been pointed out repeatedly (7,10), but these analyses are rarely implemented in electrophysiology (11). This is likely because mixed effects models take a long time to solve with iterative restricted maximum likelihood techniques, making them computationally prohibitive for large datasets. The computational cost is further exacerbated if one still wants to take advantage of nonparametric resampling and TFCE. We have developed a novel technique called Mixed Effects for Large Datasets (MELD) that makes it easy to apply permutation testing, TFCE, and LMER to the type of large datasets commonly found in neuroimaging research. Here we explain in detail how MELD works and compare its performance to the standard approach of t-tests with TFCE and permutation testing (t+TFCE).

### Linear Mixed Effects Regression

In both a typical approach and in LMER, one investigates the relationship between independent variables (such as stimulus type or reaction time) and neural data (dependent variables) at each feature, where each time-point at each electrode is a feature. The typical method for applying a general linear model to EEG data would be to fit the model to each feature within each subject and then to do a t-test on each term across subjects to determine which terms are significantly different from zero across subjects. In LMER, a single model is fit to the data from all trials and all subjects simultaneously. This is achieved by defining some model terms as “fixed effects,” or invariant across subjects, and other terms as “random effects,” or variable across subjects or trials or any other grouping factor. In a mixed effects model, the random effects are treated as if they are drawn from a distribution (the shape of that distribution can be specified, but most often it is assumed to be Gaussian).

The strengths of LMER come from the fact that it can simultaneously fit a model across multiple levels of the data, and thereby account for multiple sources of variance within a single framework. This allows LMER to perform well in unbalanced designs. LMER can be more sensitive than individual GLMs combined across subjects because in LMER, the fit of the model for each subject is informed by the models for all of the other subjects (7). This is particularly true in situations where there are low numbers of subjects or low numbers of trials per subject. A general complication of using a mixed effects model is that, due to the complex error structure, it is unclear what the degrees of freedom should be for testing the significance of each coefficient. There are different approaches to estimating the degrees of freedom that allow the significance of each coefficient to be assessed (12–14), but these methods make assumptions that may not be always be supported. Permutation testing offers a more robust alternative for determining the significance of coefficients, but permutation testing is made impractical by another weakness of LMER. Since LMERs are solved with an iterative restricted maximum likelihood approach, it can take a long time to fit a complex mixed effects model. This is particularly problematic when applying LMER to each element or feature of a neural dataset as a massive univariate approach, and then further compounded if using permutations. Consequently, LMER can be orders of magnitude slower than the tests typically used in a massive univariate approach, such as an ANOVA, t-test, or GLM. The computational cost of element-wise LMERs is such that they are infeasible for most neural datasets without use of a supercomputer, massive downsampling, or both.

## MELD: Mixed Effects for Large Datasets

Our goal for MELD was to make it possible to analyze neural data or other large datasets with LMER without a supercomputer. Again, as with a GLM or LMER, we are predicting values of the dependent neural data (e.g., combinations of time-point and electrode across events) with independent variables (e.g., experimental condition or manipulation). The primary obstacle for this is the time required to run an LMER for each dimension (or feature) of the dependent data, which we circumvent by running the LMER on the dependent data after it has been transformed to a reduced space defined via a singular value decomposition (SVD) of the correlation between the independent and dependent data. Taking the SVD of the correlation matrix identifies components that maximize the covariance between the independent and dependent variables. We then take a linear combination of dependent features weighted by each SVD component, which turns MELD into multivariate, rather than a univariate, method, increasing power while allowing us to run one LMER per component instead of one LMER per feature. Prior to the SVD, we select elements of the correlation matrix that are stable across subjects by means of bootstrapping and TFCE to improve the resilience of the SVD in the face of the large numbers of noise elements that are often present in neural datasets. Since we are running many fewer LMERs, the use of permutation testing becomes computationally feasible, and to ensure that our feature selection step has not biassed our results, we repeat the same feature selection procedure for each permutation. In MELD, the strengths of permutation, TFCE, SVD, and LMER mitigate each other’s weaknesses, producing a method that is more powerful and more versatile than the standard approach to neural data analysis.

### MELD in Detail

MELD consists of the following steps:

1. Correlate the independent and dependent data.
2. Apply TFCE.
3. Identify and mask unstable feature correlations.
4. Calculate the SVD.
5. Reduce the dimensionality of the dependent data.

a. Dot product of dependent data with right singular vectors.
6. Run LMER.

a. Predict the reduced dependent data with the independent data.
b. Transform t-values back to the feature space
7. Permutation test to assess feature level significance.

a. Dependent data is reshuffled within subjects.
b. Repeat steps **1** through **6** for each permutation.

For the sake of demonstration, we imagined a simplified experiment in which a 3 electrode x 5 timepoint set of neural data was collected in each of 10 trials for three subjects. Different items were presented in each trial so that no subject saw the same item twice. Trials were evenly split between two behavioral conditions, A and B, and there is a continuous variable with a different random value in each trial as well as mock neural data corresponding to each trial (Fig. 1A). We predict the neural data with the following mixed effects model:

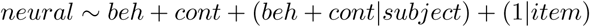

**Figure 1:**
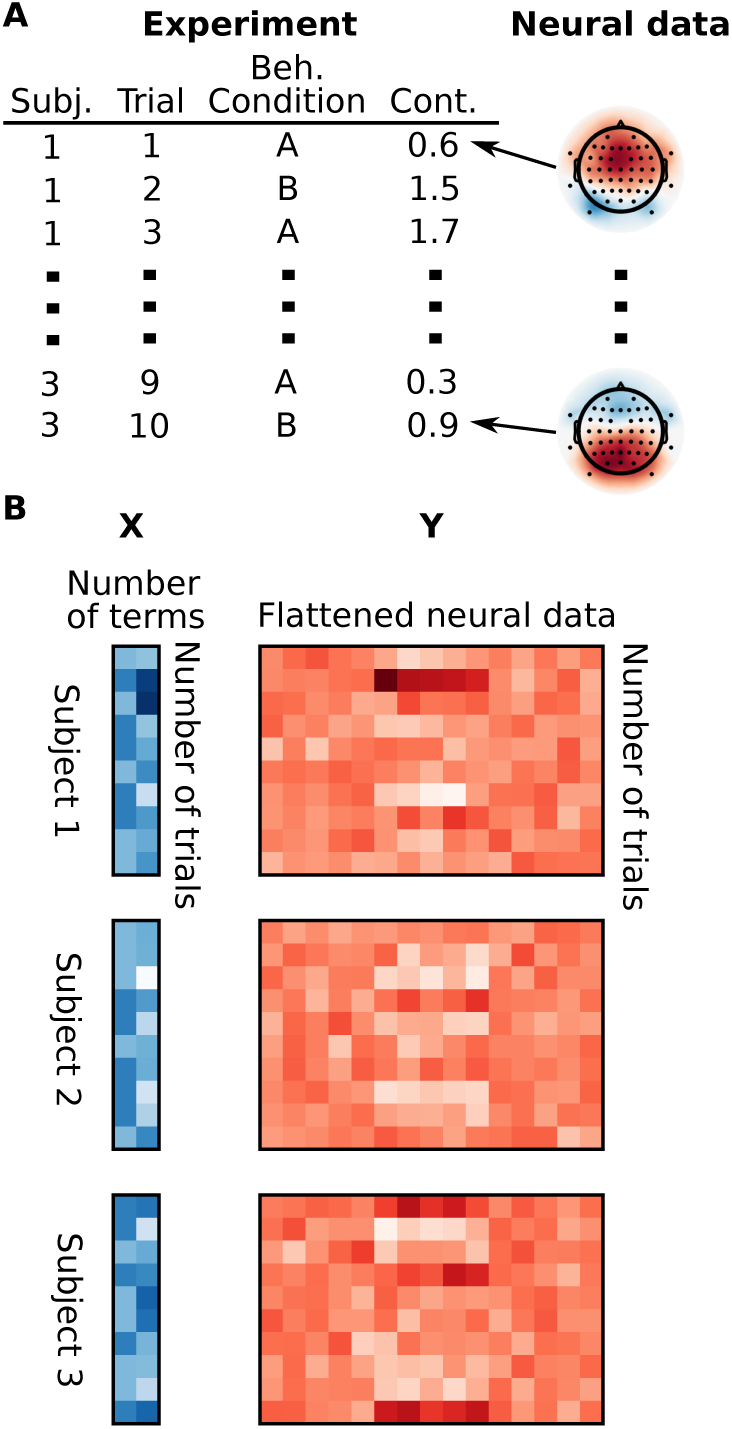
Organization of data for MELD analysis. In (A) you can see the data from a simulated experiment with two independent variables, a behavioral condition (Beh. Condition) and a continuous variable (Cont.), for each of 10 trials for three subjects and examples of corresponding mock neural dependent data. This information is assembled into matrices *X* (independent data) and *Y* (dependent data) for each subject and the matrices for each subject are stacked (B).

This model indicates that we are fitting fixed effects representing the mean effects of the behavioral condition and the continuous variable, as well as allowing the magnitude of these effects and their intercepts to vary across subjects. The intercept is additionally allowed to vary across individual items. Careful consideration should be given to the specification of the random effects. We refer readers to (15) and (7) for discussion of random effects specification. In the following description of our technique we provide the dimensions of matrices our analysis would produce based on this small dataset.

The trial information is collected in a matrix *X* with a number of rows equal to the number of trials per subject (10 trials per subject = 10 rows per subject) and number of columns equal to the number of independent variables (2 independent variables = 2 columns). The neural data are flattened and stacked to produce the matrix *Y* with number of rows equal to the number of trials per subject (10 trials per subject = 10 rows per subject) and number of columns equal to the number of features in the neural data (15 features = 15 columns) (Fig. 1**B**).

#### Step 1: Correlate the independent and dependent data

In order to calculate the correlation between the independent and dependent variables, the columns of matrices *X* and *Y* are normalized within subject so that the mean of each column is 0 and the sum of the squared values of the column is 1. The dot product of 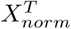 and *Y_norm_* for each subject then produces correlations in the matrix *R,* which has number of rows equal to the number of independent variables per subject (2 independent variables * 3 subjects = 6 rows) and number of columns equal to the number of features in the neural data (15 features = 15 columns).

#### Step 2: Apply TFCE

TFCE improves sensitivity for contiguous signals (in both space and time), so we enhance the values in *R* prior to feature selection in **Step 3.** First, we convert the correlation coefficients in *R* to *z*s with Fisher’s z-transformation to normalize the distribution. Then we apply TFCE to each row of *R* based on the connectivity of the features in features space, producing cluster enhanced matrix *R_e_* with the same dimensions as *R.*

#### Step 3: Identify and mask unstable feature correlations

Neural datasets typically have a large proportion of features where the correlation between dependent data and any given dimension of the independent data is not stable across subjects. The goal of this step is to remove these unstable feature correlations to improve the performance of the SVD in **Step 4.** Taking the SVD of the feature correlations identifies the transformation that reduces the dimensionality of the dependent data while maintaining the relationships between the dependent and independent data. If we did not mask unstable feature correlations, the SVD would give each unstable feature correlation very little weight, but neural datasets have so many unstable feature correlations that their cumulative effect would drown out the signal in our transformed data. It is important to note that we are determining the across subject stability of feature correlations between the dependent data and each of the independent variables separately. Feature correlations that have an unstable relationship with one predictor may be stably related to another predictor. We implement the feature correlation selection step as an across subject t-test on each element in *R_e_* versus 0. Since the values of *R_e_* may not be normally distributed after applying TFCE, we estimate the standard error as the standard deviation of a bootstrap distribution resampling subjects with replacement (16). In the work presented here we used a feature correlation selection threshold of p < 0.05 with 1000 bootstrap samples. Elements of *R_e_* for which we fail to reject the null hypothesis are not stable across subjects. It should be noted that it is not necessary to correct for multiple comparisons at this feature-selection step because we are simply assessing feature stability at this point. Feature significance is assessed at a later stage and fully corrected for multiple comparisons. This feature selection step is repeated in each permutation to ensure that the evaluation of significance accounts for this feature selection step. These elements are set to zero to produce the matrix of stable feature correlations, *R_s_,* with number of rows equal to the number of independent variables times the number of subjects (2 independent variables * 3 subjects = 6 rows) and number of columns equal to the number of features in the neural data (15 features = 15 columns).

#### Step 4: Calculate the SVD

Next we perform compact SVD on the combined *R_s_* matrix to identify the set of mutually orthogonal components that describe the variance of the stable feature correlations. In a compact SVD, only the components that describe variance in the original matrix are retained, so the SVD will generate a number of components equal to at most the number of independent variables times the number of subjects (2 independent variables * 3 subjects = at most 6 components). In our case the matrix *R_s_* is decomposed into left singular vectors *U,* singular values in diagonal matrix *S,* and right singular values *V^T^*. In MELD, *V^T^* defines the weights with which we reduce the dimensionality of our dependent data **(Step 5**); each row of *V^T^* corresponds to a component (6 components = 6 rows) and each column corresponds to a feature (15 features = 15 components). The singular values in *S* are proportional to the amount of variance explained by each component.

#### Step 5: Reduce the dimensionality of the dependent data

After calculating the SVD, we take the dot product of *V* with the unnormalized data *Y* for each subject, producing a weighted data matrix *Y_w_* with one row for each trial from each subject (3 subjects *10 trials =30 rows) and one column for each component (6 components = 6 columns). In our simulated experiment, this is a relatively modest reduction in dimensionality, from 15 original features to 6 components, but the number of components is not dependent on the number of original features, so we would have at most 6 components even if there were 100,000 features as might be found in an EEG or functional magnetic resonance imaging (fMRI) dataset. This reduction in the number of LMERs to calculate is one of the key advantages of MELD.

#### Step 6: Run LMER

We then fit a separate LMER for each component, producing *T_s_*,with a t-value for each fixed effect (2 fixed effects = 2 columns) for each component (6 components = 6 rows). We normalize the values of *S* by dividing by the sum of *S* giving *S_norm_*, which represents the proportion of variance accounted for by each component, and then we transform *T_s_* to feature space as follows:

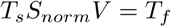

#### Step 7: Permutation test to assess feature level significance

Once the above process has been completed for the true data, we repeat the process, complete with feature selection, for permutations in which the dependent data is shuffled relative to the independent data within subjects. We apply the sum of *S* from the true data to calculate *S_norm_* for each permutation so that the feature level t-values in each permutation are scaled relative to the amount of variance explained in the true data. Monte Carlo sampling of the permutation distribution produces a distribution of t-values for each term at each feature. The distribution of t-values for each term may be scaled differently, so we create a term-level null distribution from the most extreme (greatest absolute value) t-value across features, for each permutation. With these term-level null distributions we compute empirical p-values for each term at each feature. Using the most extreme t-value from each permutation allows us to control for multiple comparisons across features (3). Once we have a p-value for each term at each feature and permutation, we then generate an across-term null distribution as the minimum p-value across terms and features for each permutation. This distribution is then used to derive the final empirical p-values for each feature in the true data, fully accounting for multiple comparisons across features and terms.

### Benefits of MELD

We expect that MELD will have several advantages over alternative approaches. Unlike t+TFCE, the sensitivity and precision of MELD will be less affected by the spatiotemporal distribution of signal features because the SVD is a multivariate technique that can combine the contribution of non-contiguous features, allowing improved sensitivity for noncontiguous signals over t+TFCE. MELD incorporates mixed effects modeling, simultaneously accounting for subject and trial level random effects. Accounting for more sources of known variance will also contribute to greater sensitivity for MELD compared to t+TFCE. Lastly, we expect that MELD will have similar computational requirements as t+TFCE. We will evaluate the performance of MELD in comparison to massive univariate approaches in the form of t-tests with and without TFCE on simulated data and on ERP data from a standard item-recognition experiment.

## Methods

We conducted two experiments using simulated data to evaluate the performance of MELD in comparison to t-tests at each element with a Monte Carlo permutation test and t-tests at each element followed by TFCE with a Monte Carlo permutation test. Our simulated data contains a binary signal with added Gaussian noise. Although smoothness is an important feature of real neural signals, smoothing the data would create ambiguity about which features constitute true positives, necessitating the definition of a cutoff and complicating assessment of each method’s performance (5). We then compared the performance of MELD and t-tests with TFCE on real EEG data.

### Simulation Experiment A

In the first simulation experiment we used a 100 x 100 field of noise with a standard normal distribution with 100 features containing signal in one of three patterns: central, split, or dispersed (Fig. 2). These signals will benefit from TFCE to varying degrees, which we will explore in greater detail in Simulation Experiment B.

**Figure 2:**
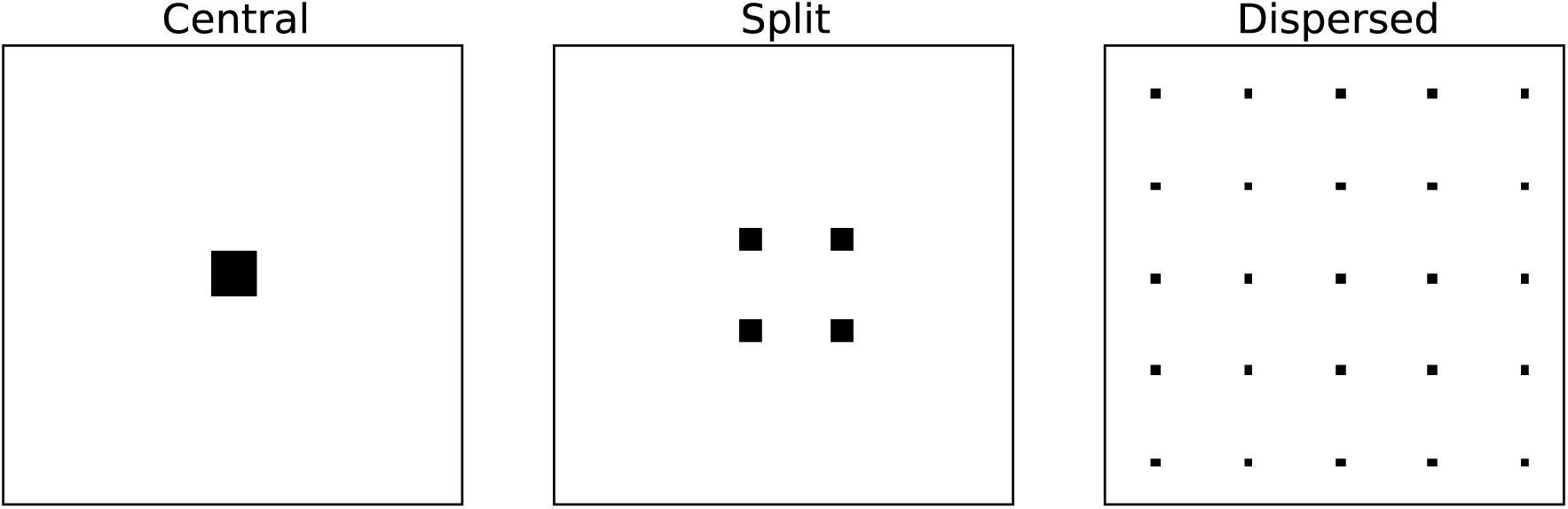
Patterns of signal used in Simulation Experiment A. These are the patterns of signal that were added to 100 x 100 noisy fields to generate simulated neural data. Central is a single central 10 x 10 cluster of 100 features. Split has four separate 5 x 5 clusters, each containing 25 features. Dispersed has 25 separate 2 x 2 clusters, each containing 4 features.

We generated signal with the following model:

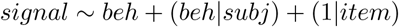

The slope of the effect for the categorical variable “beh” was varied from 0.1 to 0.9 in steps of 0.2. For “beh” category A this slope was multiplied by 0.5 and for category B it was multiplied by -0.5, producing the contrast that the three methods will be trying to detect. The item level variance on the intercept was drawn from a standard normal distribution. The subject level variance on the intercept was drawn from a normal distribution with a mean of zero and a standard deviation of 0.1. The subject level variance on the slope was also drawn from a normal distribution with a mean of zero and a standard deviation of 0.1. We simulated data for an experiment in which 9 subjects completed 50 total trials of two different types (A and B). The signal for each trial was multiplied by the signal pattern and embedded in a field of noise drawn from a standard normal distribution. In each simulation, we applied all three methods to each simulated dataset so that we could compare their performance. At each level of slope and pattern of signal we ran 20 different simulations, for 300 total simulations, to thoroughly assess the performance of the methods we are considering. Additionally we ran 100 simulations with each signal pattern with no fixed effect and the same levels of subject and item level noise to verify that the family-wise error rate (FWER; explained in **Performance Metrics** below) was adequately controlled.

To perform the t-tests we averaged the trial level signals for each of the two behavioral conditions within subjects and then ran a paired t-test comparing the signal associated with each behavioral condition across subjects. For t+TFCE, this was followed by TFCE with dt = 0.05, E = 2/3, H = 2.0, the optimal parameters as suggested by Smith & Nichols (2009). For both t-test methods we ran 500 permutations and used the most extreme value from each permutation for the null distribution.

For the MELD analysis we fit the following model:

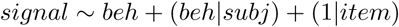

This is the same as the model used to generate the data, which we chose because the ability to account for known sources of variance is one of the advantages of LMER and we wanted to include this effect in our evaluation of MELD in comparison to standard techniques which cannot account for it.

For all three approaches we used an alpha of 0.05. For MELD we used a feature correlation stability threshold of 0.05.

This method of simulating data allowed us to have complete control over the parameters of the signal being detected. If we had applied smoothing to the data, the identity of true positive and true negative voxels would be ambiguous; this approach for generating simulated data allows us to rigorously assess the performance characteristics of the methods we are comparing. Similarly, we chose not to use simulated ERP data because of the arbitrary cutoff required to distinguish signal and noise voxels, which introduces unnecessary error into the assessment of performance of the different techniques (5).

### Simulation Experiment B

For the second simulation experiment, we were interested in comparing the performances of MELD and t-tests (with and without TFCE) at various levels of signal for signal patterns with varying degrees of centrality versus dispersion. In order to do this we ran simulations similar to those in experiment A, except that the patterns of signal represented different proportions of the central and dispersed data patterns. In all of these patterns we retained the same overall size of the field (100 x 100 features) and the same number of features containing signal (100, or 1%). In describing these patterns of signal, we refer to the percent centrality, which we define as the percentage of features in the largest cluster. The central pattern from Simulation Experiment A represents 100% centrality, and the dispersed pattern from experiment A represents 4% centrality (Fig. 3).

**Figure 3:**
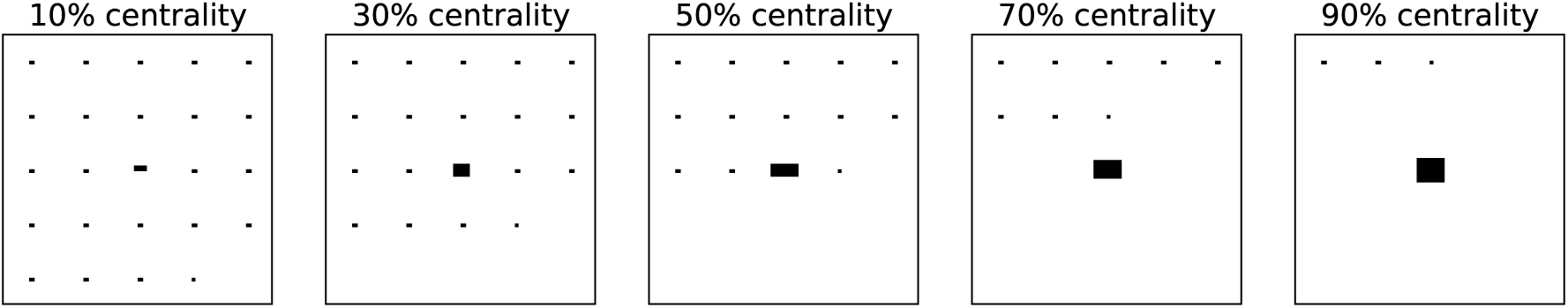
Patterns of signal used in Simulation Experiment B. These are the patterns of signal that were added to 100 x 100 noisey fields to generate simulated neural data for Simulation Experiment B. Percent centrality is defined as the percentage of features in the largest cluster.

For Simulation Experiment B we applied the same methods for all three techniques as in Simulation Experiment A.

### Performance Metrics

In evaluating performance on our simulated data we considered three metrics at a feature level: sensitivity, positive predictive value (PPV; also referred to as precision), and Matthew’s correlation coefficient (MCC). Alpha was set to 0.05 for all three methods, as in (5). Examining performance at an alpha of 0.05 instead of testing multiple thresholds to calculate an ROC is more representative of the way that these thresholds are employed in neuroimaging research. Sensitivity is equivalent to the true positive rate (TPR), the ratio of features correctly identified as containing signal to the number of features actually containing signal, and can be thought of as the ability of a technique to detect a true signal. PPV is the ratio of features correctly identified as containing signal to the total number of features identified as containing signal. PPV can be thought of as the confidence that a feature identified as containing signal actually contains that signal. The PPV is also one minus the false discovery rate. Sensitivity and PPV are important in the analysis of neural data because it is often the case that relatively few features of a neural dataset will contain signal so an ideal technique will identify all of the features with signal and as few false positives as possible. Lastly we include MCC as a metric to summarize the overall performance of each method (17). The MCC is given by the following equation:

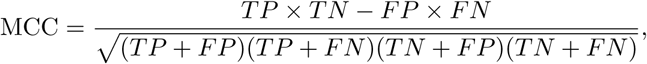

where TP is true positive, TN is true negative, FP is false positive, FN is false negative. MCC is essentially the correlation between the true signal and the signal found by a given method. The key advantage of MCC for our purposes is that it is not biased in situations such as this in which few features contain signal. In addition to the feature level performance metrics we also examine the FWER. When performing a statistical test at multiple features, the FWER is the probability of having at least one false positive across all of the features examined. Permutation based testing as implemented in MELD should have a FWER that is not significantly different from alpha (3).

### Statistical analysis

For both simulation experiments, we compared performance on all three feature level metrics aggregating across all of the simulations with signal. We evaluated MELD versus t+TFCE and MELD versus t-test. For both of these comparisons, we tested the null hypotheses that TPR, MCC, and PPV were equivalent between methods using a bootstrap t-test. We tested the null hypothesis that each method’s FWER was equal to alpha in the absence of a fixed effect with an exact binomial test. Results were visualized using the Python packages Matplotlib (18) and Seaborn (19).

### EEG Data Collection

We examined data from an experiment using a standard item recognition procedure in which a lists of words are presented followed by test lists containing both studied and non-studied words. For every test word, subjects had to decide whether it was studied or not.

#### Subjects

Twenty-four right-handed subjects (14 female) between 18 and 38 years of age (M = 21.6) were recruited from the university community at The Ohio State University. All subjects spoke and read English fluently. Subjects provided written consent in accordance with requirements of the local institutional review board and were paid $20 for their time. Data from subjects for whom there were excessive motion artifacts or recording noise (n = 5) or who failed to follow instructions (n =2) are not included in the analyses. For the analysis presented here we randomly selected 10 subjects from the 17 with good data. A different, but overlapping, subset of data from this experiment was previously used to validate a method for topographic latent source analysis (20) and a larger subset of data from this experiment was used for a single trial analysis of EEG in recognition memory (21).

#### Stimuli

Stimuli were drawn from the SUBTLEXUS database (22). A total of 549 high-frequency words (between 40 and 400 occurrences per million, M=122.31, SD=84.52) and 553 low-frequency words (between 1 and 7 occurrences per million, M=1.44, SD = 0.39) formed the stimulus pool. All words in the pool had between 4 and 7 letters, and word length was equated across the frequencies (M=5.03 and 4.99, respectively).

#### Design

The experiment used a 2 (Word Frequency: high vs. low) x 3 (Item Strength: strong vs. weak vs. new) factorial within subject and within-list design. Each subject studied and was tested on 13 lists of words. Each study list was constructed by randomly selecting 9 words of each frequency from the pool to be studied one time (the “weak” condition) and 9 words of each frequency to be studied three times (the “strong” condition). Thus, there were 36 unique words and 72 item presentations in each study list, with the order of the presentations pseudo-randomized such that there were no immediate repetitions of the strong items. A matching test list consisting of all 36 studied words along with a set of 36 matching lures was constructed for each study list. The order of words on the test lists was randomized in 2 blocks with the first 18 unique studied words tested in the first block and the second 18 tested in the second block. This was done to ensure that the end of list items were never tested immediately, thereby reducing the probability of recency effects. After discounting the first list, which was used for practice, this design yielded a total of 108 target trials and 108 lure trials for each experimental condition. This study was carried out in accordance with the recommendations and approval of the Ohio State University Social and Behavioral Sciences Institutional Review Board with written informed consent from all subjects. All subjects gave written informed consent in accordance with the Declaration of Helsinki.

#### Equipment

A desktop computer with a 17” LCD display was used to present the stimuli and collect subject responses. A custom program written using the Python Experiment Programming Library (PyEPL) (23) was used to generate the study lists for each subject, control the timing of the tasks, and record subject responses. Scalp EEG data were sampled and recorded reference-free at 1000 Hz using a DC-powered actiCHamp amplifier/analog-to-digital converter connected to a desktop PC equipped with PyCorder software (BrainProducts GmbH, Munich, Germany). Prior to receiving instructions, each subject was fitted with an elastic cap containing 64 active electrodes arrayed in an extended 10-10 layout. Electrode impedances were reduced to less than 25K ohms in accordance with operating instructions for the actiCAP system (BrainProducts GmbH, Munich, Germany).

#### Procedure

After informed consent was obtained, subjects were fitted with the EEG cap, seated in front of a computer, and given instructions by the experimenter. Subjects were told that they would be studying lists of words and would be given a recognition memory test after each study list. They were informed that some of the study words would be repeated but were not told that this was an experimental manipulation. Subjects were also instructed to try to avoid thinking back to previous words on the list during the study session. Subjects were then given the first study and test list as a practice list. The experimenter answered any questions and then started the experiment proper.

Prior to each study list, an orientation cross was displayed in the center of the screen for 2600-3000 ms. The study list words were then displayed, one at a time, in the center of the screen for 1600 ms followed by a blank screen for 300-700 ms. Following the study list, an orientation cross was displayed for 1000-1200 ms to signify the test was about to begin. On each test trial, the probe word was displayed at the center of the screen and a prompt “Old or New?” was displayed below it. Subjects made an old judgment by pressing the “J” key on a standard keyboard with the index finger of their right hand or a new judgment by pressing the “K” key with the middle finger of their right hand. Once the subject made a response, the prompt was removed from the screen. Subjects were allowed up to 1800 ms to make a response, and then the stimulus was removed from the screen. Test trials were separated by a jittered inter-trial interval of 400-800 ms.

### EEG Data Processing

All preprocessing was done using the Python Time Series Analysis (ptsa) library (https://github.com/compmem/ptsa). Voltage measurements from each electrode were first re-referenced to the average of the right and left mathematically-linked mastoid electrodes at each time-point then high-pass filtered at 0.5 Hz. Eye-blinks and minor motion artifacts were corrected using the wavelet-enhanced independent components analysis algorithm (24). Following eye blink correction, the voltage data from the test phase were segmented into events using a 1.25 s window starting 0.25 s prior to the onset of each stimulus. Any epoch that exhibited a kurtosis value greater than 5.0 or a voltage range exceeding 100 microvolts was removed from the analysis. ERPs were calculated separately for each channel and down-sampled to 100 Hz for subsequent analyses. This eyeblink correction algorithm cleans eyeblinks from the EEG signal quite well, but it does not completely remove artifacts due to horizontal eye-movements, which were removed with the Gratton method (25).

### EEG data analysis

We analyzed the EEG data for the contrast between the perceived recall and viewing a subjectively novel item. This is one form of the classic “Old/New” paradigm (26). Specifically, this analysis compares the ERP for trials in which participants responded “New” and trials in which they responded “Old.” Our analysis focused on the period between 0.1 s and 1 s after stimulus onset. For the t-test based analyses, evoked time courses for each type of trial were averaged within subjects and the t-test was conducted across subjects. For MELD, we fit the following model:

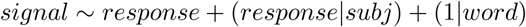

The signal was the trial level evoked potential, 90 time points at 64 channels for 5760 features, the stimuli for each subject were drawn from the same pool, and the term “(1|word)” indicates that we are accounting for fluctuations in the signal due to the word presented.

## Results

We conducted two experiments using simulated data to evaluate the performance of MELD in comparison to t-tests at each element with a Monte Carlo permutation test (referred to as t-tests) and t-tests at each element followed by TFCE with a Monte Carlo permutation test (referred to as t+TFCE). In the presence of both trial level and subject level noise, derivation of a signal-to-noise ratio is complex, so we kept the noise contribution constant while varying the magnitude, or slope, of the effect of the behavioral condition. The data in our simulation experiments consisted of a simple two dimensional field with different shapes of signal. We did not apply any smoothing to the signal because this would create ambiguity about which features should be counted as true positives. In other words, this allowed us to avoid having to define a threshold amount of signal required to be present in a feature for it to be considered a signal-containing feature.

In Simulation Experiment A we evaluated MELD, t+TFCE, and t-test alone with different patterns of signal and different amounts of signal. Simulation Experiment A indicated that MELD and t-test with TFCE exhibited different sensitivities to signal distribution, so we systematically varied the signal distribution in Simulation Experiment B to better characterize this effect. We then analyzed real experimental data from an item recognition task with MELD and t+TFCE and t-tests to demonstrate the feasibility of using MELD with real world data.

### Simulation Experiment A

The goal of Simulation Experiment A was to get an initial understanding of the performance of MELD in comparison to typical approaches under a variety of experimental conditions. We evaluated the performance of MELD and t-tests between conditions with and without TFCE in simulated data with three different patterns of signal and 5 different levels of contrast between conditions. The different levels of contrast correspond to different slopes in the model used to generate the signal.

#### Timing

We ran 20 simulations at each combination of settings for a total of 300 runs. All the simulations were carried out on compute nodes in the Oakley cluster at the Ohio Supercomputing Center. Each node has a 12 core Intel Xeon x5650 with 48 GB of ram. The average CPU time for MELD in this experiment was 352.70 minutes (0.27 SEM). The average CPU time for calculating the t-test was 7.37 minutes (0.01 SEM). The average CPU time for t+TFCE was 193.30 minutes (0.11 SEM). Assuming cores with equivalent performance, we can estimate that the compute times on an 8 core desktop would be 1 minute for the t-test, 24 minutes for t+TFCE, and 44 minutes for MELD. MELD does take longer to run than t+TFCE, but the compute time is on the same order of magnitude, and well within the capabilities of a modern desktop or a higher end laptop.

#### Aggregate performance

Generally an investigator will not have prior knowledge of the characteristics of the true signal in a dataset. In order to replicate this situation in judging MELD, we assessed aggregate performance across all 300 of the simulations with signal from experiment A (Fig. 4). MCC measures the overall agreement between a true signal and the result of a statistical test. Based on MCC (Fig. 4A), MELD performed better overall than t+TFCE (t-value = 6.42; p-value < 0.001) or t-test (t-value = 18.20; p-value < 0.001). MELD had better overall performance than t+TFCE in 191 simulations (63.7% of 300) and better overall performance than t-test in 222 simulations (74.0%). Furthermore, there were only 13 simulations (4.3% of 300) in which t+TFCE had a greater MCC than MELD, and only 2 (0.7%) in which t-test had a better MCC than MELD.

**Figure 4:**
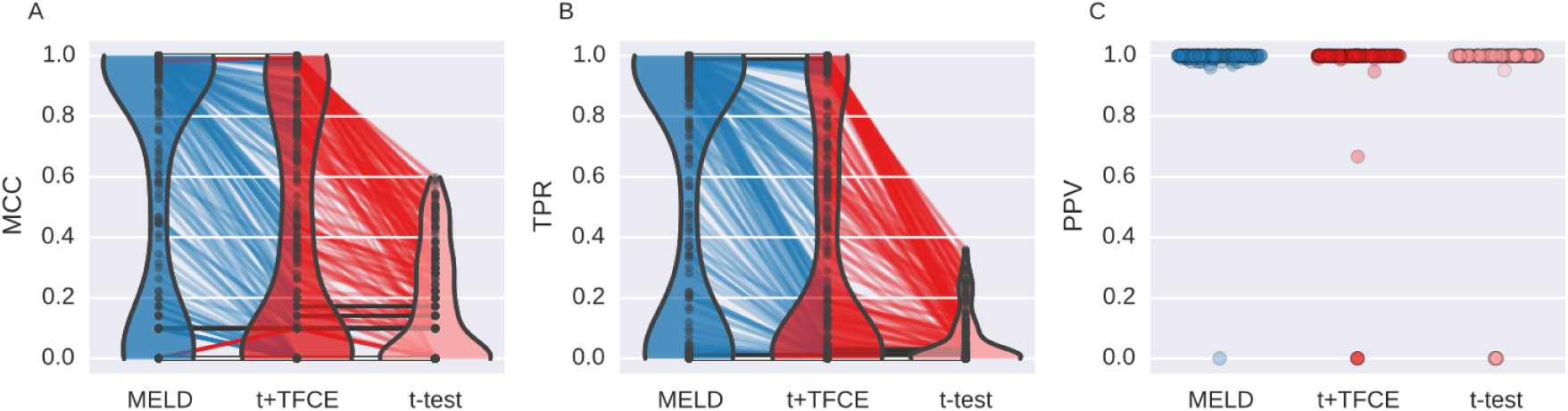
Aggregated results of Simulation Experiment A. This shows the performance of the three different methods (MELD, t+TFCER, and t-test) using three metrics (MCC,TPR, and PPV) for each of 300 simulations. Violin plots in A and B demonstrate the distributions of results for MCC and TPR respectively. The width of the colored section indicates the kernel density estimate. MELD has a greater density of values with high MCC and TPR than either of the two other methods. The colored lines connecting the points indicate results from the same simulation; lines are colored based on the method for which they had the highest value. Gray lines indicate equal values. The paucity of red lines between MELD and t+TFCE reflects very low number of simulations for which t+TFCE performed better than MELD. The distribution of PPV (C) was divided almost entirely between 0 and 1, resulting in a misleading violin plot, so instead we have just plotted a point for each simulation result. The width over which points are spread is indicative of the number of points present at that value. PPV is undefined when a method fails to identify any features as containing signal, so there are unequal numbers of points for each method.

Improved overall performance for MELD seems to be the result of increased sensitivity (Fig. 4B).

MELD had a greater TPR than t+TFCE (t-value = 5.66; p-value < 0.001) or t-test (t-value = 15.63; p-value < 0.001). MELD had a higher sensitivity than t+TFCE in 188 simulations (62.7%) and a higher sensitivity than t-test in 219 simulations (73.0%). t+TFCE only had a higher sensitivity than MELD in 2 simulations (0.7%), and t-test only had a higher sensitivity in 1 (0.3%). In practice, being able to accurately detect signal at more features is desirable, but even more important is whether or not you identify any signal at all. There were 41 simulations (13.6%) in which MELD was the only technique that identified signal, finding an average of 13.6 (2.52 SEM) features (out of 100) in these cases. In only one simulation (0.3%) t+TFCE and t-test identified a single feature while MELD failed to find anything.

As reflected in the greater overall performance, MELD’s greater sensitivity does not come at the cost of a reduction in PPV (Figure 4C). In Simulation Experiment A, there was not sufficient evidence to reject the null hypothesis that MELD had the same PPV as t+TFCE (t-value = 1.48; p-value = 0.16), nor was there sufficient evidence to reject the null hypothesis that MELD had the same PPV as t-test (t-value = 1.96; p-value = 0.07; note this is a marginal effect where MELD has a better PPV than t-test).

The expanded results of Simulation Experiment A reiterate the aggregated results (Fig. 5) with MELD clearly outperforming t+TFCE and t-test across the full range of simulations. Because of the complex noise structure used to simulate the data, traditional metrics of signal to noise ratio are difficult to calculate, so we instead refer to the slope of the ‘beh’ fixed effect used to simulate the 0 data. At a slope of 0.1, none of the methods performed well. At a slope of 0.9, MELD and t+TFCE have excellent performance, particularly for the central signal pattern. MELD and t+TFCE both outperform t-test at all except the lowest level of signal.

**Figure 5:**
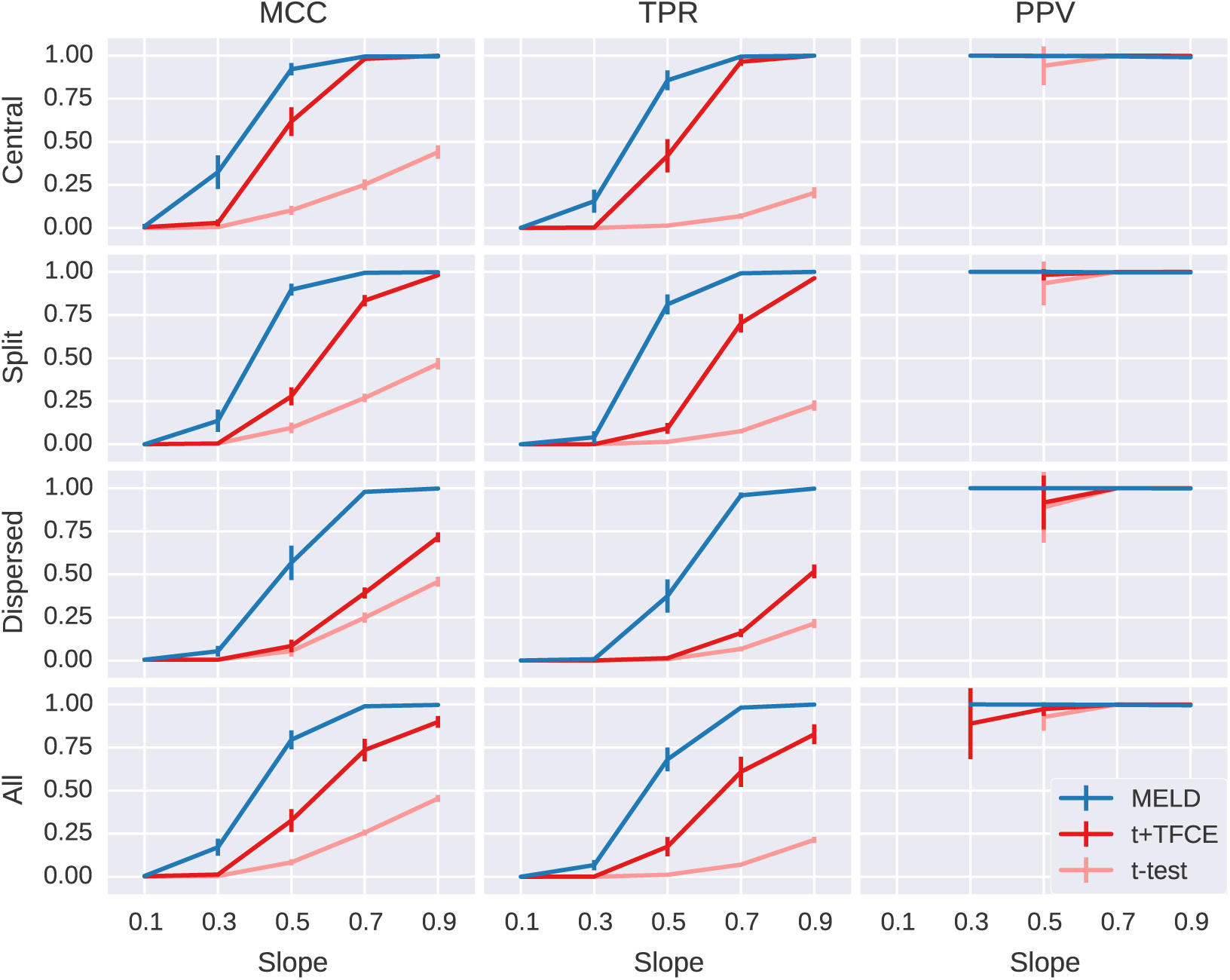
Expanded results of Simulation Experiment A. This shows the performance of the three different methods (MELD, t+TFCE, and t-test) on simulated data using three metrics (MCC, TPR, and PPV). Each row shows the results for one signal pattern and the final row shows the results collapsed across all signal patterns. Slope refers to the level of contrast between conditions; higher values indicate a stronger signal. Error bars indicate the 95% confidence interval across the simulations run at those conditions. PPV is undefined if a method does not find any significant elements; consequently, PPV results are unreliable for points at which fewer than 5 simulations identified significant features, so these points are not shown. MELD has the clearest advantage for lower levels of signal and for the dispersed signal pattern.

As expected, since t-test does not incorporate information about the signal distribution, it is unaffected by the signal pattern, while signal pattern has the strongest effect on t+TFCE. MELD is somewhat affected by the signal pattern, but to a much lesser degree than t+TFCE, giving MELD the greatest advantage for the distributed signal pattern. We explore the effect of signal dispersion on the performance of MELD and t+TFCE in greater detail in Simulation Experiment B.

Additionally, we ran 300 simulations with no fixed effect to assess the FWER of these methods. The alpha for all three methods was set to 0.05. Over 300 simulations with no fixed effect, MELD had false positives on 16 out of 300 simulations (5.33%). Based on the binomial test there is not sufficient evidence to reject the null hypothesis that MELD’s FWER is different from 0.05 (p = 0.79). There was not sufficient evidence to reject this null hypothesis for t+TFCE or t-test either; these methods each found false positives in 10 simulations (3.33%; p = 0.231).

## Simulation Experiment B

In Simulation Experiment A it appeared that the “clustered-ness” or centrality of the signal had a stronger impact on t+TFCE than on MELD. We systematically investigated the differential effects of signal distribution on the performance of MELD, t+TFCE in Simulation Experiment B with simulated data across 5 different levels of centrality and at five different slopes. In terms of the relative performance of MELD, t+TFCE, and t-test, Simulation Experiment B replicates the results of experiment A (Fig. 6). MELD had a higher MCC than t+TFCE (t-value = 8.04; p-value < 0.001) or t-test (t-value = 20.20; p-value < 0.001). MELD was more sensitive than t+TFCE (t-value = 9.75; p-value < 0.001) or t-test (t-value = 23.30; p-value < 0.001). Finally, there was not sufficient evidence to reject the null hypothesis that MELD had the same PPV as t+TFCE (t-value = 0.57; p = 0.58) or t-test (t-value = 0.15, p = 0.85).

**Figure 6:**
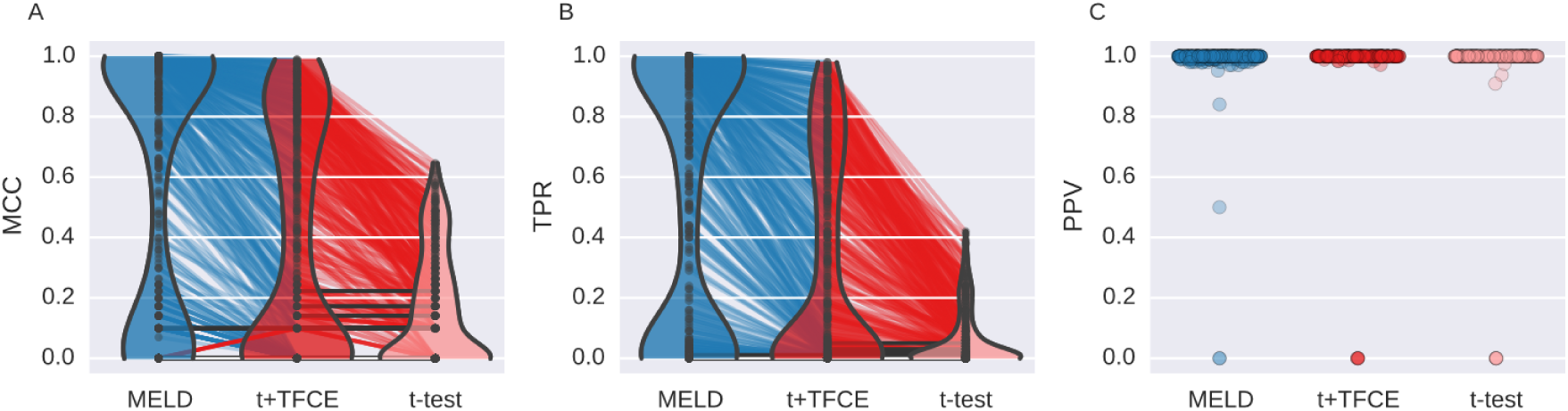
Aggregated results of Simulation Experiment B. This shows the performance of the three different methods (MELD, t+TFCER, and t-test) using three metrics (MCC,TPR, and PPV) for each of 500 simulations. Violin plots in A and B demonstrate the distributions of results for MCC and TPR respectively. The width of the colored section indicates the density of points. MELD has a greater density of values with high MCC and TPR than either of the two other methods. The colored lines connecting the points indicate results from the same simulation; lines are colored based on the method for which they had the highest value. Gray lines indicate equal values. The paucity of red lines between MELD and t+TFCE reflects very low number of simulations for which t+TFCE performed better than MELD. The distribution of PPV (C) was divided almost entirely between 0 and 1, resulting in a misleading violin plot, so instead we have just plotted a point for each simulation result. The width over which points are spread is indicative of the number of points present at that value. PPV is undefined when a method fails to identify any features as containing signal, so we were unable to connect points across simulations.

The interaction of slope, centrality, and analysis method are shown in Figure 7. In our simulations, t-test alone did not appear to be affected by the centrality of signal, only by its intensity (represented by slope). t+TFCE was strongly affected by the centrality of the signal, with performance dropping to that of the t-test with less central (more dispersed) signals. This demonstrates that TFCE improves sensitivity for signals with larger extents in space or time, but not for isolated signals, as expected. At slopes of 0.7 and 0.9 MELD detected all the signal regardless of centrality and at slopes of 0.1 and 0.3 t+TFCE detected very little signal, so the results from simulations with slope of 0.5 provide the best window into the effects of signal distribution on the performance of MELD. In particular, comparing the impact of centrality on the performance of t+TFCE with the impact of centrality on the performance of MELD at a slope of 0.5 allows us to infer how much MELD’s performance benefits from the use of TFCE as part of the feature selection step 2.

**Figure 7:**
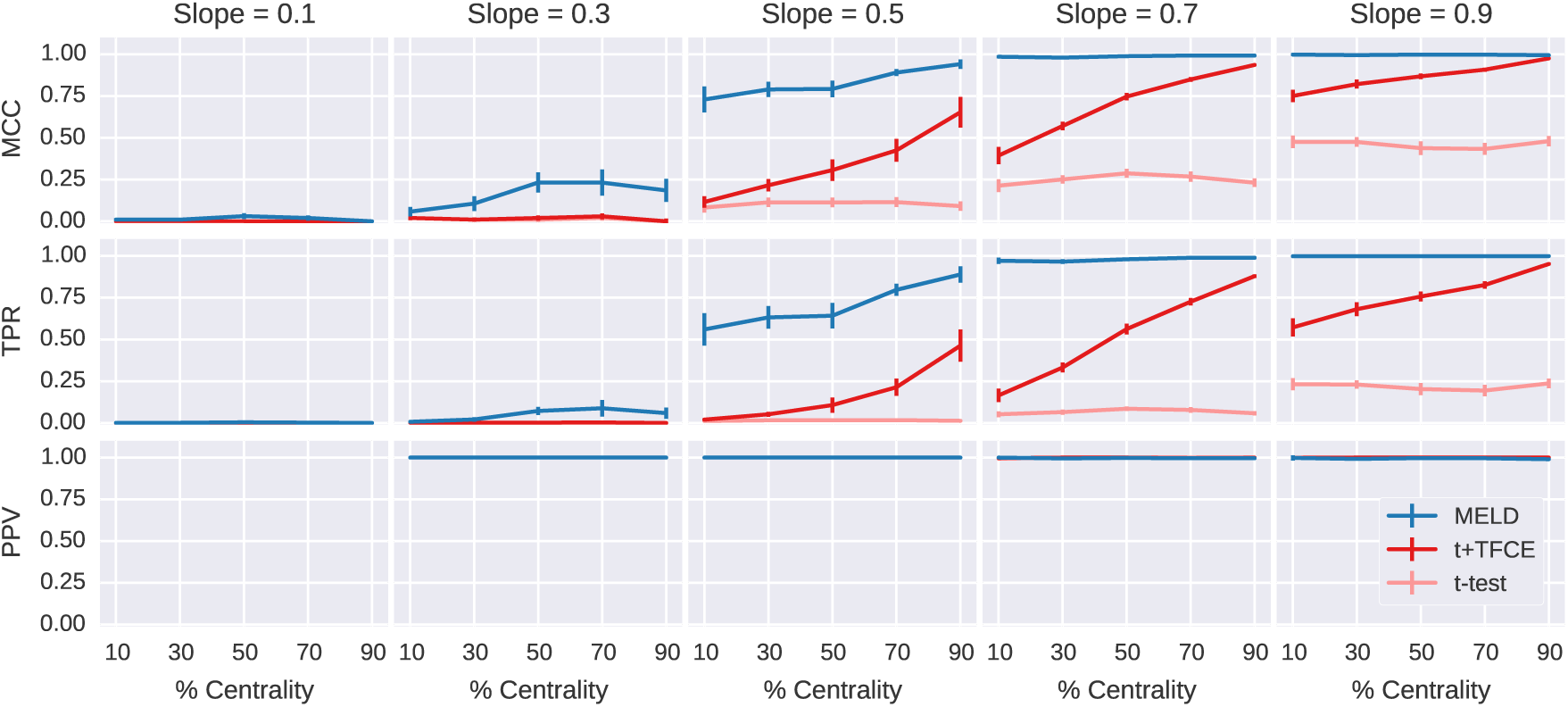
Detailed results of Simulation Experiment B. The results for MCC, TPR, and PPV from 500 simulations at 5 different slopes and 5 different centralities demonstrate the reduced impact of centrality on MELD when compared to t+TFCE. t-test was unaffected by centrality but was much less sensitive than the other two methods. MELD was stable across all levels of centrality at higher levels of slope while t+TFCE was sensitive to differences in centrality at all levels of slope. At a slope of 0.5, MELD was more sensitive to changes in centrality than at higher slopes, but at 10% centrality it still has an MCC of 0.72 ± 0.08 95%CI, compared to t+TFCE with an MCC of 0.12 ± 0.03 95%CI. The gain in overall performance and sensitivity for MELD is clearest at a slope of 0.5, but importantly, MELD is also able to detect some signal at a slope of 0.3, when the other methods are detecting very little.

If we assume that the relationship between MELD with and without TFCE would be similar to that of t-tests with and without TFCE, then the sensitivity of MELD at a slope of 0.5 and a centrality of 10% (0.56 ± 0.10 95%CI) approximates MELD’s sensitivity without TFCE. We compared this to MELD’s sensitivity at slope of 0.5 and a centrality of 90% (0.89 ± 0.05 95% CI) to estimate that TFCE contributed roughly 37% of MELD’s sensitivity at a slope of 0.5 and centrality of 90%. At the same level of slope and centrality, t+TFCE had a sensitivity of 0.46 ± 0.10 95% CI, while t-test had a sensitivity of 0.013 ± 0.005 95% CI, thus TFCE contributed 71.2% of the sensitivity of t+TFCE at a slope of 0.5 and centrality of 90%. This rough quantification indicates that TFCE accounts for a smaller portion of MELD’s sensitivity than it does for t+TFCE. Overall, MELD showed strong performance in Simulation Experiment B, and the interactions between slope, centrality, and performance demonstrate the contribution of TFCE to MELD’s power.

### MELD analysis of EEG data

We performed an analysis of EEG data from an item recognition task with MELD, t-test, and t+TFCE to demonstrate the feasibility of using MELD with real world data. In this item recognition task, participants were presented with words and asked to indicate whether or not they recognized them from a previous study period. We compared the difference in evoked response between words that participants thought were new to words that they thought had been presented previously (a standard old/new recognition memory comparison). The t-test alone was not able to identify stable features, so we focus here on the performance of t+TFCE and MELD. Both methods were able to identify a strong posterior old/new effect. However, MELD identified a signal with greater extent in time (Fig. 8) and across the scalp surface (Fig. 9) that was more similar to the canonical LPC seen across the literature (26–30) than significant features identified to be more posterior LPC by the t+TFCE method. The results from MELD are also accounting for random effects at the subject and item level. It may be due to these random effects, which are difficult to display graphically, that MELD is not finding a few timepoints around 0.7 s significant while t+TFCE is.

**Figure 8:**
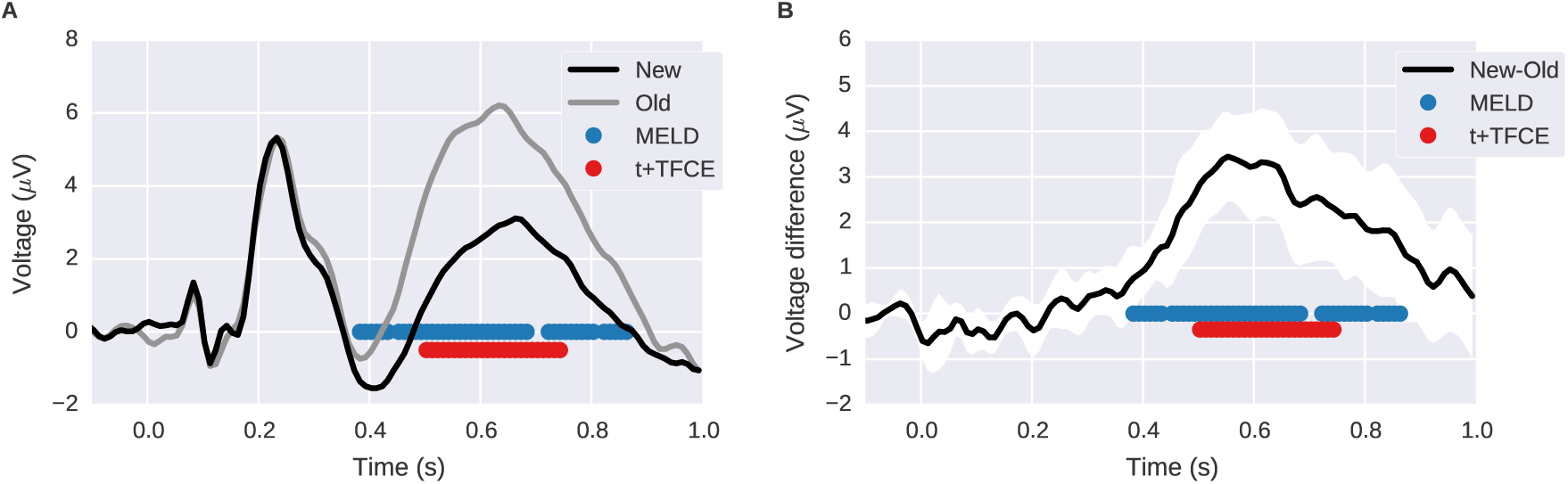
Time course of old/new effect assessed by MELD and t+TFCE. A shows the grand average ERP at channel Pz for old and new trials. B shows the within subject mean difference with the 95% confidence interval in white around it. The blue and red dots indicate time points identified as significant by MELD and t+TFCE respectively.

**Figure 9:**
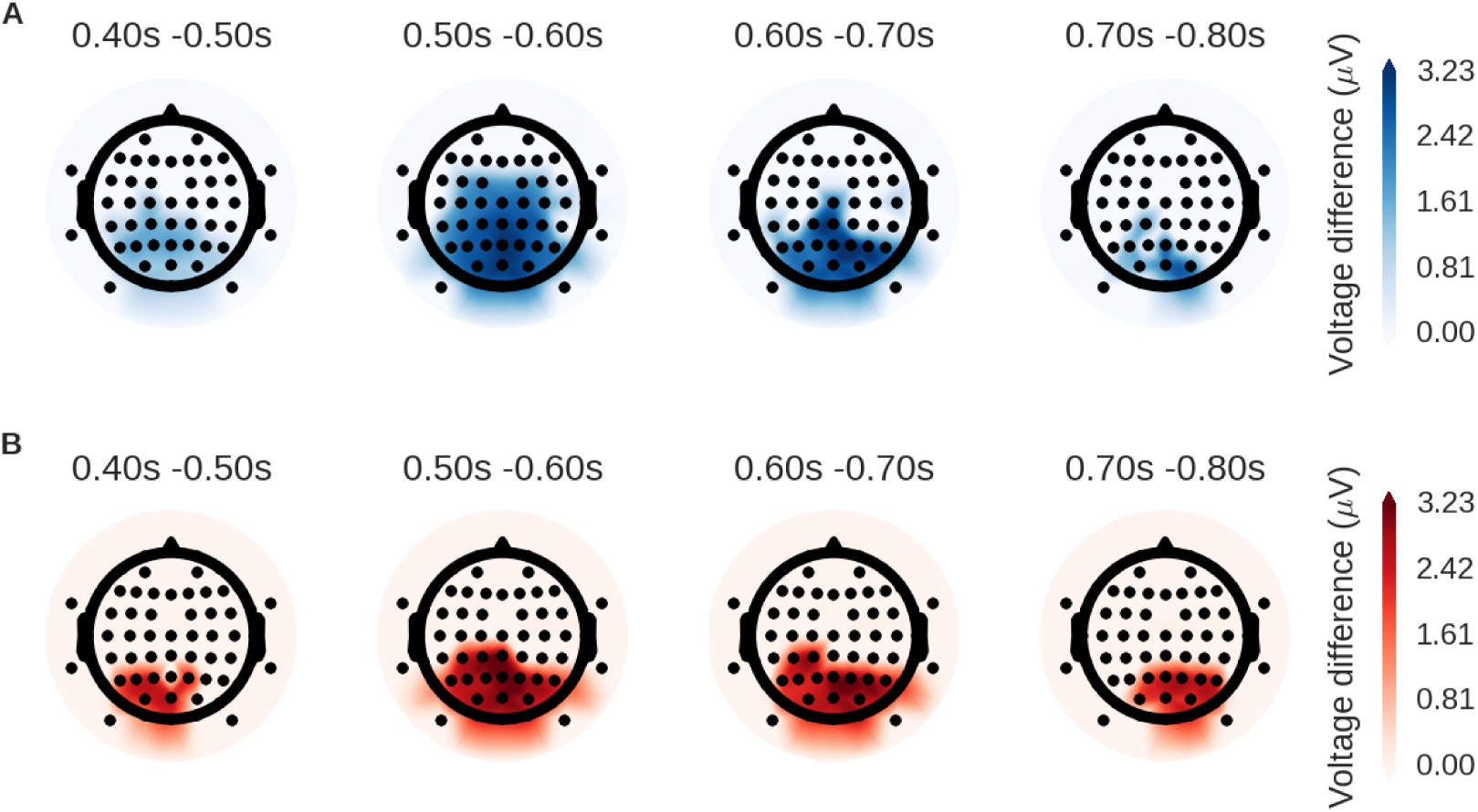
Scalp distribution of old/new effect assessed by MELD and t+TFCE. These topographic plots represent a top down view of the head with the triangle at the top representing the nose. Each dot indicates the location of an EEG channel. Average differences in voltage for significant sensor-channel combinations are plotted for both MELD (A; in blue) and t+TFCE (B; in red). Color only signifies the method used, not the direction of the effect, which is positive for both MELD and t+TFCE. Both techniques identify similar regions, but the effect identified by MELD has greater extent in time and space.

## Discussion

MELD is a novel multivariate approach that enables efficient application of mixed effects models to large datasets. We first evaluated MELD’s performance on simulated data so that we could accurately compare it to the most popular massive univariate techniques for EEG analysis: t-tests comparing conditions with and without TFCE tested via permutation test.

In Simulation Experiment A, MELD consistently demonstrated better overall performance and greater sensitivity than either univariate method without a significant difference in PPV across the range of conditions we tested (Fig. 4-7). Our results for t+TFCE are similar to those obtained in another simulation study, which found that TFCE performed very well, but was somewhat conservative (5). MELD is somewhat more computationally intensive than t+TFCE, but the improved performance more than compensates for the longer computation time.

MELD continued to demonstrate better performance than the mass univariate approaches in Simulation Experiment B when we varied the amount of clustering of the signal features. As expected, t-tests with TFCE were sensitive to the distribution of signal features, showing the strongest performance on highly clustered data (Fig. 7). MELD’s performance was negatively affected by increased dispersion of signal features, but to a lesser extent than t+TFCE. The results of Simulation Experiment B make it clear that MELD is more powerful than the current standard technique across a broad range of conditions because of the combined contributions of the SVD and TFCE.

Unlike our simulation experiments where we generated both the signal and the noise, in our EEG experiment (and any actual real-world neural experiment) it is impossible to know the “ground truth” for the neural data against which to compare the results from our different approaches with different sample sizes or different numbers of trials. That said, the results from the EEG experiment do support the conclusions from our simulation results. MELD identified a significant late positive component in central, parietal, and occipital electrodes that was more similar to the canonical LPC seen across the literature (26–30) than the more posterior late positive component identified by the t+TFCE method. It also identified a larger region of significance, both in time and space, further suggesting an ability to extract signal more efficiently from noise. In the future, we plan to utilize larger neural datasets, including those of different modalities, to further characterize the performance of MELD.

This is the initial publication of MELD, and the technique does have some limitations. Currently the only method for dealing with missing data is to either drop that feature from all subjects, or to drop that trial. In the future we hope to implement a method for estimating missing data in order to make MELD more flexible. MELD has one free parameter: the threshold for the initial feature selection step. A full exploration of the interaction of this parameter with characteristics of the data is outside of the scope of this report, but in our experience the value we use here, 0.05, works well. The ultimate feature level p-value is determined by a permutation test, and the value for this threshold will always be consistent across permutations, so while it may impact the sensitivity of the technique, varying the feature selection threshold is unlikely to increase the FWER. In the future, as MELD is validated on new types of data – such as MEG and fMRI data – we will investigate the effects of this parameter further.

With the use of bootstrapping, permutation testing, and SVD, MELD may seem at first to be quite similar to partial least squares (PLS). PLS also uses these techniques, but the approach, goals, and interpretation of PLS are quite different. PLS is a multivariate linear correlation technique that can identify the relationship between multiple, collinear independent variables and dependent variables (31). In PLS the results of the SVD are interpreted as orthogonal latent variables. The significance of each latent variable is assessed by permutation testing. Bootstrap resampling across subjects then identifies which features are stably contributing to each significant latent variable. In MELD, identification of stable features with bootstrap resampling and TFCE is done before the SVD, and the SVD is carried out on cluster enhanced correlations. The SVD in MELD serves to reduce the dimensionality of the neural data, which allows us to employ an LMER simultaneously modeling variance at subject and trial level for each component. In PLS, the SVD necessarily collapses across trials within subjects, preventing assessment of trial level random effects. In MELD, the t-values from the LMER are mapped back to feature space, and the permutations then allow significant features to be identified while controlling for multiple comparisons across features and terms. The goal of PLS is the identification of distinct components, and it is the significance of these components that is assessed by permutation. While “non-rotated” PLS can be used for hypothesis testing, the most typical use of PLS is as an exploratory analysis to identify latent relationships between the dependent and independent data, leaving interpretation of significant components up to the researcher. MELD is explicitly designed to facilitate testing a specific hypothesis about the relationship between independent and dependent data formulated as an LMER model.

MELD could also be compared to feature reduction methods such as ICA, but these approaches are unsupervised and not guaranteed to identify the variance that separates conditions of interest. In contrast, the feature reduction step in MELD specifically selects features with stable differences across the conditions of interest. In ICA, interpretation of the identified components for the research questions of interest is often complicated and can invite post-hoc explanations. MELD, however, is designed to test specific hypotheses about the the relationship between the neural data and conditions of interest, while controlling for random effects at multiple levels. Thus, even though they both provide multivariate solutions to dimensionality reduction, ICA and MELD represent two distinct analysis approaches, with MELD performing both dimensionality reduction and hypothesis testing.

In this initial presentation of MELD we have focussed on analysis of EEG data from a straightforward experimental design, but MELD is much more flexible than this. In our examples we have had random effects at the subject and item level, but MELD could also be applied to a within subject analysis with random effects at the block and item level, or subject, block, and item level effects could be considered simultaneously. Beyond neural data, MELD is applicable in any situation in which you would like to use a mixed effects model and have too many features for element-wise LMER to be feasible. While we have focused on MELD with TFCE, it is possible to run MELD without TFCE for datasets in which connectivity between features is difficult to define, such as genetics or economics.

In summary, we have demonstrated that MELD is a much more powerful technique than those that are currently used. We hope that the ability to efficiently apply mixed effects analyses to large datasets will facilitate the use of novel study designs to address hypotheses previously beyond the reach of typical data analysis approaches in neuroscience and beyond.

## Acknowledgements

This work was supported in part by an allocation of computing time from the Ohio Supercomputer Center. D.M.N. was supported by the Choose Ohio First for Bioinformatics Scholarship.

## References

1. RossionB, CaharelS.ERP evidence for the speed of face categorization in the human brain: Disentangling the contribution of low-level visual cues from face perception. Vision Research. 2011 Jun; 51(12):1297–311.

2. DegabrieleR, LagopoulosJ, MalhiG.Neural correlates of emotional face processing in bipolar disorder: An event-related potential study. Journal of Affective Disorders. 2011 Sep;133(1):212–20.

3. MarisE, OostenveldR.Nonparametric statistical testing of EEG- and MEG-data. Journal of Neuroscience Methods. 2007 Aug;164(1):177–90.

4. Smith SM, Nichols TE.Threshold-free cluster enhancement: Addressing problems of smoothing, threshold dependence and localisation in cluster inference. NeuroImage. 2009 Jan;44(1):83–98.

5. MensenA, KhatamiR.Advanced EEG analysis using threshold-free cluster-enhancement and non-parametric statistics. NeuroImage. 2013 Feb;67:111–8.

6. PernetCR, LatinusM, NicholsTE, RousseletGA. Cluster-based computational methods for mass univariate analyses of event-related brain potentials/fields: A simulation study. Journal of Neuroscience Methods. 2014;

7. BaayenRH, DavidsonDJ, BatesDM.Mixed-effects modeling with crossed random effects for subjects and items. Journal of Memory and Language. 2008 Nov;59(4):390–412.

8. BeckmannCF, JenkinsonM, SmithSM.General multilevel linear modeling for group analysis in FMRI. NeuroImage. 2003 Oct;20(2):1052–63.

9. CnaanA, LairdNM, SlasorP.Using the general linear mixed model to analyse unbalanced repeated measures and longitudinal data. Statistics in Medicine. 1997 Oct;16(20):2349–80.

10. BagiellaE, SloanRP, HeitjanDF.Mixed-effects models in psychophysiology. Psychophysiology. 2000 Jan;37(1):13–20.

11. Amsel BD.Tracking real-time neural activation of conceptual knowledge using single-trial event-related potentials. Neuropsychologia. 2011 Apr;49(5):970–83.

12. KenwardMG, RogerJH.Small sample inference for fixed effects from restricted maximum likelihood. Biometrics. 1997 Sep;53(3):983–97.

13. Satterthwaite FE.An Approximate Distribution of Estimates of Variance Components. Biometrics Bulletin. 1946 Dec;2(6):110–4.

14. Welch BL.The Generalization of “Student’s” Problem when Several Different Population Variances are Involved. Biometrika. 1947 Jan;34(1/2):28–35.

15. BarrDJ, LevyR, ScheepersC, TilyHJ.Random effects structure for confirmatory hypothesis testing: Keep it maximal. Journal of memory and language. 2013;68(3):255–78.

16. EfronB, TibshiraniRJ.An Introduction to the Bootstrap. CRC Press; 1994.

17. Matthews BW.Comparison of the predicted and observed secondary structure of T4 phage lysozyme. Biochimica et Biophysica Acta (BBA) - Protein Structure. 1975 Oct;405(2):442–51.

18. Hunter JD.Matplotlib: A 2D graphics environment. Computing In Science & Engineering. 2007;9(3):90–5.

19. WaskomM, BotvinnikO, HobsonP, ColeJB, HalchenkoY, HoyerS, et al.Seaborn: V0.5.0 (November 2014). 2014.

20. GershmanSJ, BleiDM, NormanKA, SederbergPB.Decomposing spatiotemporal brain patterns into topographic latent sources. NeuroImage. 2014 Sep;98:91–102.

21. RatcliffR, SederbergPB, SmithTA, ChildersR.A single trial analysis of EEG in recognition memory: Tracking the neural correlates of memory strength. Neuropsychologia.2016 Dec;93(Pt A):128–41.

22. BrysbaertM, New B.Moving beyond Kucera and Francis: A critical evaluation of current word frequency norms and the introduction of a new and improved word frequency measure for American English. Behavior Research Methods. 2009 Nov;41(4):977–90.

23. GellerAS, SchleiferIK, SederbergPB, JacobsJ, KahanaMJ.PyEPL: A cross-platform experiment-programming library. Behavior research methods. 2007 Nov;39(4):950–8.

24. CastellanosNP, MakarovVA.Recovering EEG brain signals: Artifact suppression with wavelet enhanced independent component analysis. Journal of Neuroscience Methods. 2006 Dec;158(2):300–12.

25. GrattonG, ColesMG, DonchinE.A new method for off-line removal of ocular artifact. Electroencephalography and Clinical Neurophysiology. 1983 Apr;55(4):468–84.

26. RuggMD, CurranT.Event-related potentials and recognition memory. Trends in Cognitive Sciences. 2007 Jun;11(6):251–7.

27. CurranT.Effects of attention and confidence on the hypothesized ERP correlates of recollection and familiarity. Neuropsychologia. 2004;42(8):1088–106.

28. DüzelE, YonelinasAP, MangunGR, HeinzeH-J, TulvingE.Event-related brain potential 5 correlates of two states of conscious awareness in memory. Proceedings of the National Academy of Sciences. 1997 May;94(11):5973–8.

29. Smith ME.Neurophysiological Manifestations of Recollective Experience during Recognition Memory Judgments. Journal of Cognitive Neuroscience. 1993;5(1):1–13.

30. StrózakP, BirdCW, CorbyK, FrishkoffG, CurranT.FN400 and LPC memory effects for 0 concrete and abstract words. Psychophysiology. 2016 Nov;53(11):1669–78.

31. KrishnanA, WilliamsLJ, McIntoshAR, AbdiH.Partial Least Squares (PLS) methods for neuroimaging: A tutorial and review. NeuroImage.2011 May;56(2):455–75.

